# Azithromycin robustly synergizes with tetracyclines to overcome acquired minocycline resistance in *Acinetobacter baumannii*

**DOI:** 10.64898/2026.01.14.699369

**Authors:** Syed Hasan Raza, Suman Tiwari, Nicholas Dillon

## Abstract

Multidrug-resistant (MDR) *Acinetobacter baumannii* is a critical nosocomial pathogen with rapidly shrinking treatment options, and although minocycline (MIN) remains useful, resistance is increasingly common, highlighting the need for effective combination therapies. Here, we investigate azithromycin (AZM)-tetracycline synergism as a strategy to overcome acquired MIN resistance and define the antibiotic-specific drivers of this interaction. Using MIC_90_ and checkerboard fractional inhibitory concentration (FIC) assays, we tested AZM and MIN across multiple A. baumannii strains (AB5075, ATCC19606, BAA1710, BAA1789) in both bacteriological (CA-MHB) and physiologic (RPMI+) media and found AZM-MIN synergy to be robust across strains and media, with enhanced synergy in RPMI+. To determine whether this effect reflects a class-wide phenomenon, we systematically evaluated different macrolide-tetracycline combinations and observed that synergy occurred consistently in AZM containing pairs, identifying AZM (not tetracycline selection) as the dominant driver of synergy. Mechanistically, translation inhibition assays using an AB5075-luxCDABE reporter demonstrated that AZM uniquely sustained strong translational inhibition at multiple sub-inhibitory fractions, supporting a kinetic basis for synergy when combined with tetracyclines. Finally, AZM-MIN synergy was evaluated against 41 MDR isolates from the CDC-FDA Antimicrobial Resistance Isolate Bank, where synergism was detected in 8/41 strains and reduced the inhibitory MIN concentration below the CLSI resistance breakpoint for 4 isolates, indicating that AZM-based combinations can partially restore MIN susceptibility in resistant backgrounds. Together, these findings establish AZM as a potent and distinctive macrolide synergist with tetracyclines against MDR A. baumannii and support further exploration of AZM-tetracycline combination therapy to extend the clinical utility of MIN and mitigate selection for resistance.

## Introduction

The global rise of infections with multidrug-resistant (MDR) bacterial pathogens are one of the greatest threats to modern medicine. Among these MDR organisms, the Gram-negative coccobacillus *Acinetobacter baumannii* is especially concerning owing to its extensive genomic plasticity, ability to persist in hospital settings, and capacity to rapidly acquire antibiotic resistance (1, 2). Alarmingly, MDR strains of *A. baumannii* are already estimated to account for 20.3% of hospital-acquired infections across WHO-defined regions in Europe, the Eastern Mediterranean, and Africa (3). Successful treatment of *A. baumannii* infections has become increasingly difficult, often necessitating the use of second-line agents such as polymyxins and tetracycline derivatives (4). Yet, the pathogen’s ability to rapidly acquire new resistance mechanisms continues to outpace available therapies, underscoring the urgent need for novel treatment strategies.

MIN has emerged as a particularly promising option for treating highly resistant *A. baumannii* infections. MIN is a second-generation tetracycline that binds to the 30S ribosomal subunit and inhibits protein synthesis by preventing the entry of transfer RNA into the aminoacyl site, effectively halting peptide chain elongation (5). Unlike other tetracyclines, MIN retains efficacy against many MDR *A. baumannii* strains, even those harboring common the TetA efflux pump which confers resistance to doxycycline and tetracycline (6). This resistance evasion is partly due to MIN’s increased lipophilicity and steric hindrance, allowing it to bypass TetA efflux systems (7). The treatment efficacy of MIN is supported by a 2025 retrospective study of patients with carbapenem-resistant *A. baumannii* pneumonia, which reported higher treatment success rates in patients receiving MIN (58.3%) either as monotherapy or in combination with colistin, TGC, or ampicillin/sulbactam, compared to those who did not receive MIN (41.7%) (8). Furthermore, MIN susceptibility remains prevalent in *A. baumannii* as a 2024 cross-sectional study examining 100 clinical *A. baumannii* isolates found that 50 of 83 (60.2%) MDR strains were susceptible to MIN (9). These findings underscore the continued in vitro activity of MIN against *A. baumannii*. Taken together with clinical outcomes data, they support the essential role of MIN as an effective therapeutic option for MDR *A. baumannii* infections.

Beyond its individual efficacy, we previously demonstrated that MIN exhibits unexpected synergism when combined with the macrolide AZM against MDR *A. baumannii* strain AB5075 (10). Although both antibiotics target the bacterial ribosome and were not predicted to interact synergistically, our study showed that each agent complemented kinetic deficiencies of the other i.e. MIN providing rapid but short-lived inhibition of translation, and AZM supplying delayed yet sustained inhibition. This complementary action resulted in potent synergy in both bacteriological and physiological media, as confirmed by checkerboard assays, bacterial cytological profiling, and translation activity assays. In murine pneumonia models, AZM-MIN therapy significantly reduced lung bacterial burden and improved survival compared to either drug alone. Importantly, this synergy extended beyond *A. baumannii* to other high-priority MDR pathogens such as *Klebsiella pneumoniae, Pseudomonas aeruginosa*, and methicillin-resistant *Staphylococcus aureus* (10). These findings established that AZM can act as a powerful partner to MIN by stabilizing and prolonging its activity, allowing the use of lower concentrations of both antibiotics while still achieving robust bacterial killing. By broadening the therapeutic window of MIN and extending its utility to otherwise resistant isolates, our earlier work provided the foundation for deeper exploration into the drivers of AZM-mediated synergism and its potential to be generalized across other macrolide-tetracycline pairings.

Despite these promising findings, key questions remained unresolved. It was unclear whether the observed synergy was unique to MIN or represented a generalizable feature of AZM across the tetracycline class. Moreover, while our earlier work highlighted complementary inhibition kinetics, the specific contribution of AZM relative to its macrolide counterparts had not been defined. Evidence indicates that AZM may possess unique properties that enable sustained translation inhibition and drive synergistic effects, but the generality of this phenomenon across different antibiotics, strains, and media conditions is still unresolved(10-12). Furthermore, although AZM is widely prescribed and has a well-established safety profile, its unconventional activity against Gram-negative pathogens remains underappreciated. These limitations highlight the need for studies that clarify AZM’s distinctive contributions to synergistic therapy and evaluate its potential for broader clinical application.

Building on our prior work, the present study directly investigates how AZM enables synergy with tetracyclines against MDR *A. baumannii*. We first assess AZM-tetracycline combinations across multiple clinical isolates to determine whether AZM driven synergy is strain dependent or broadly conserved. We then compare AZM with other macrolides to define whether this effect is specific to AZM or generalizable within the class. Using translational reporter assays, we quantify the kinetics of protein synthesis inhibition to clarify how AZM’s prolonged activity complements tetracycline action. Finally, we evaluate whether AZM can lower tetracycline inhibitory concentrations below established clinical breakpoints, effectively restoring susceptibility in resistant strains. Through these investigations, we position AZM not simply as a partner to tetracyclines but as the key driver of synergy, with the potential to expand therapeutic strategies against MDR *A. baumannii* and related pathogens.

## Methods

### Bacterial Strains

*Acinetobacter baumannii* AB5075, a model strain isolated from a patient with tibial osteomyelitis, was obtained from the Walter Reed Army Medical Center (13). The laboratory drug-sensitive strain ATCC19606, originally isolated in 1948 from a urinary tract infection, was accessioned by the American Type Culture Collection (ATCC) in 1966 (14). Two clinical MDR isolates, BAA1710 (from human blood) and BAA1789 (from a tracheal aspirate), were also obtained from ATCC (15). The AB5075-luxCDABE strain was constructed by inserting the Tn7-luxCDABE mini-Tn7 element into the attTn7 site of AB5075, as previously described (13, 16). Additionally, 41 MDR *A. baumannii* isolates were acquired from the CDC-FDA Antimicrobial Resistance Isolate Bank, a publicly available resource for researchers (17).

### Antibiotics

Powdered forms of minocycline (MIN, Melinta Therapeutics, NDC 70842-160-10), omadacycline (OMA, MedChemExpress, HY-14865), doxycycline (DOX, Fresenius Kabi, 1311), tetracycline (TET, Thermo Fisher Scientific, J61714-14), azithromycin (AZM, Athenex Pharmaceuticals, 70860-100), roxithromycin (ROX, MedChemExpress, HY-B0435), erythromycin (ERY, Sigma Aldrich, E5389), and clarithromycin (CLR, Sigma Aldrich, PHR1038) were purchased from their respective vendors.

Master stocks were prepared as follows: MIN (20 mg/mL), DOX (50 mg/mL), and AZM (100 mg/mL) were solubilized in water; OMA (100 mg/mL), TET (30 mg/mL), ROX (100 mg/mL), and ERY (50 mg/mL) were solubilized in dimethyl sulfoxide (DMSO); and CLR (10 mg/mL) was solubilized in acetone. All antibiotic stocks were stored at –80 °C and diluted in water to the desired experimental concentrations prior to use.

### Culturing conditions and media

Bacteria were streaked on Luria agar from frozen stocks (20% glycerol in CA-MHB) stored at –80 °C and inoculated into 3 mL CA-MHB in 14 mL Falcon round-bottom snap-cap tubes (Corning #352001). Cultures were incubated overnight at 37 °C with shaking at 200 RPM in a New Brunswick Innova 44/44R incubator. The following day, 30 µL of overnight culture was subcultured into biological triplicates (3 mL each) in either CA-MHB or RPMI+ media and grown until mid-log phase (OD_600_ ≈ 0.4). All experiments were conducted in Costar 96-well flat-bottom plates (Corning #3370) at a final volume of 200 µL/well.

### MIC assays

Minimum inhibitory concentrations (MIC_90_), defined as the lowest antibiotic concentration that inhibits ≥90% of bacterial growth relative to untreated controls, were determined as previously described (10, 18). In 96-well plates (Corning #3370), each well received either 20 µL of water (control) or 20 µL of a 10× antibiotic stock. Mid-log cultures were diluted to 5 × 10^5^ CFU/mL (OD_600_ ≈ 0.002), and 180 µL of this suspension was added to each well to reach a final volume of 200 µL. Plates were incubated at 37 °C and 200 RPM for 19 hours. Bacterial growth was assessed by OD_600_ readings using a Biotek Synergy H1 multimode plate reader(18).

### FIC assays

Fractional inhibitory concentration (FIC) assays were conducted to assess antibiotic synergy, following previously described methods (10). For each well, 20 µL of water (control) or 10 µL each of two 20× antibiotic stocks were added. Then, 180 µL of diluted mid-log culture (OD_600_ ≈ 0.002) was added to reach a total volume of 200 µL. Plates were incubated for 19 hours at 37 °C and 200 RPM. Growth was measured at OD600 using the Biotek Synergy H1.

FIC values were calculated by dividing the concentration of each drug in the combination by its respective MIC. The FIC index (FICI) was the sum of the two FICs. A FICI ≤ 0.5 was considered indicative of synergism (19). All FIC assays were conducted in biological triplicates. Representative FIC plots display a single replicate from each triplicate set.

### Translation inhibition assays

The AB5075-*lux*CDABE strain constitutively expresses the *lux* operon, producing both luciferase and its substrate to yield luminescence under normal translational conditions. This luminescence is reduced in the presence of translation-inhibiting antibiotics, enabling quantification of translation inhibition (10, 13, 16).

AB5075-*lux* was cultured and sub-cultured in CA-MHB using the same protocol as the MIC assays. For single-drug assays, 20 µL of a 10× antibiotic stock was added to each well of a 96-well clear-bottom plate (Thermo Fisher #165306). For dual-drug assays, 10 µL of two different 20× stocks were added per well. A total of 180 µL of diluted culture (≈5 × 10^5^ CFU/mL; OD_600_ ≈ 0.002) was added to each well. Plates were incubated at 37 °C in the Biotek Synergy H1 without shaking for 15 hours.

Luminescence and OD_600_ readings were recorded every 30 minutes. Luminescence was normalized to OD_600_ to control for bacterial density. The normalized luminescence values were then divided by untreated controls to calculate the percent translational activity.

### Statistics

All statistical analyses were conducted using GraphPad Prism 10. One-way ANOVA was used to evaluate the effects of monotherapy and combination therapy treatments. Dunnett’s multiple comparison test was applied to identify significant differences between treatment groups and controls. Statistical significance was defined as *p* ≤ 0.0500, with notations as follows: *p* value ≤0·0500 with * ≤0·0500–0·0100, ** ≤0·0100–0·0010, *** ≤0·0010– 0·0001, and **** ≤0·0001.

## Results

### Minimum Inhibitory Concentrations of AZM and MIN against MDR A. baumannii

Our prior studies indicated that AZM and MIN synergized against multiple bacterial pathogens including *A. baumannii* strain AB5075 (10). To assess if the synergy was specific to AB5075 or broadly impacted other *A. baumannii* strains we first examined how various strains responded to both AZM and MIN in CA-MHB and RPMI+. As the determination of a MIC is the gold standard for assessing the potency of an antibiotic, we first measured the MIC_90_ values of AZM and MIN in monotherapy against four strains of *A. baumannii*: MDR laboratory strain AB5075, antibiotic-sensitive laboratory strain ATCC19606, and MDR clinical isolates BAA1710 and BAA1789. MIC_90_ assays were completed in either CA-MHB, a standard rich bacteriological medium, or RPMI+, the physiological media consisting of the bicarbonate buffer system. 10% Luria Broth was added to RPMI to supplement bacterial growth (RPMI+). Across all four strains, MIN was more potent in CA-MHB (MIC_90_ 0·5 to 8 µg/ml) but drops significantly in activity in RPMI+ (MIC_90_ > 256 µg/ml) (Table 1). In contrast, AZM was shown to be ineffective against each *A. baumannii* strain in CA-MHB (64 to 128 µg/ml) but displays improved potency in RPMI+ (0·25 to 1 µg/ml) (Table 1).

**Table 1.** MIC_90_ (µg/ml) against *A. baumannii* strains in CA-MHB and RPMI+.

|  | CA-MHB |  | RPMI+ |  |
| --- | --- | --- | --- | --- |
|  | MIN | AZM | MIN | AZM |
| *Acinetobacter baumannii* |  |  |  |  |
| - AB5075 | 1 | 128 | >256 | 1 |
| - ATCC19606 | 0.5 - 1 | 128 | >256 | 1 |
| - BAA1710 | 4 | 128 | >256 | 0.5 - 1 |
| - BAA1789 | 4 - 8 | 64 | >256 | 0.25 - 0.5 |

### AZM-MIN Synergy is Strain and Conditionally Independent

We next assessed if AZM and MIN were synergistic against each *A. baumannii* strain. We utilized fractional inhibitory concentration (FIC) assays to determine the extent of AZM-MIN synergism against each strain in CA-MHB. Synergy occurs when the calculated fractional inhibitory concentration index (FICI) is ≤ 0·5. FICI values within the range >0·5 - 4 are indicative of additivity, and FICI values > 4 display an antagonistic relationship between the two drugs. AZM and MIN combinations were synergistic, with FICI values ranging from 0·25-0·5, against each strain in CA-MHB (Fig 1. a-d). We next conducted the checkerboard assays in RPMI+ to determine if the synergism was dependent on media selection. Synergy was found to be even more pronounced in RPMI+ with FICI values of 0.09-0.31, likely owing to the strongly reduced MIN activity in RPMI+ (Fig 1. e-h).

**Figure 1.**
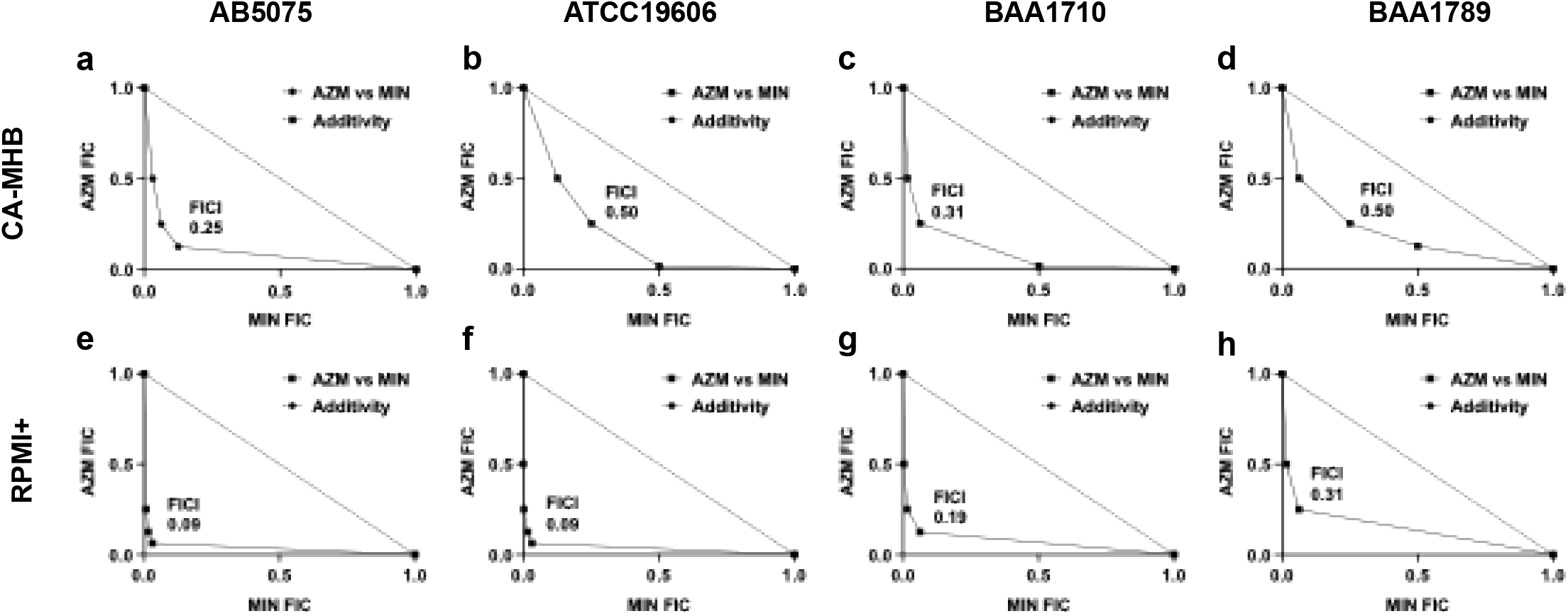
AZM + MIN combination therapy is synergistic independent of the media or *A. baumannii* strain. FIC assays of AZM + MIN combinations were performed on four different *A. baumannii* MDR strains in CA-MHB (a-d) or RPMI+ (e-h): *(a,e)* AB5075, *(b,f)* ATCC19606, *(c,g)* BAA1710, and *(d,h)* BAA1789. All experiments were completed in triplicates, with a representative plot shown. FIC values are relative to 1.0, which is set as the MIC for each antibiotic in monotherapy. A synergistic relationship between the antibiotics is indicated when FICI ≤ 0.5.

### AZM Drives Synergism Across Macrolide-Tetracycline Combinations

Prior studies with other antibiotic classes, like β-lactams and aminoglycosides, have suggested synergistic interactions can be a class wide phenomenon (20). Having established AZM-MIN synergism occurs independently of strain and media selection, we next sought to determine if synergy was specific between AZM and MIN, or broadly between macrolides and tetracyclines. Three different antibiotics from each of the macrolide (ROX, ERY, and CLR) and tetracycline (OMA, DOX, and TET) drug classes were selected for additional FIC analysis to assess if synergy was a class wide phenomenon. MIC_90_ assays were first conducted for each drug in CA-MHB against AB5075 to establish the appropriate ranges for the FIC analysis (Table 2). Similarly, FIC assays were performed in identical conditions and organized such that each macrolide was tested in combination with each tetracycline. Of the 16 total FIC assays, synergy was observed in all combinations that contained AZM (FICI 0·25-0·5) (Fig 2 a-d), as well as the OMA + ERY (FICI 0·31) (Fig. 2j) combination. All other combinations demonstrated an additive relationship (FICI >0·5). Of the synergistic interactions, AZM-MIN proved to be the most effective with a FICI = 0·25 (Fig. 2a). These results indicate AZM, and not tetracycline selection, is a key driver of synergy.

**Table 2.** MIC_90_ (µg/ml) of macrolide & tetracycline antibiotics against AB5075 in CA-MHB.

|  | Macrolides |  |  |  | Tetracyclines |  |  |  |
| --- | --- | --- | --- | --- | --- | --- | --- | --- |
|  | AZM | ROX | ERY | CLR | MIN | OMA | DOX | TET |
| - AB5075 | 128 | 64 | 32 | 32 | 1 | 2 | 2 | 4 |

**Figure 2.**
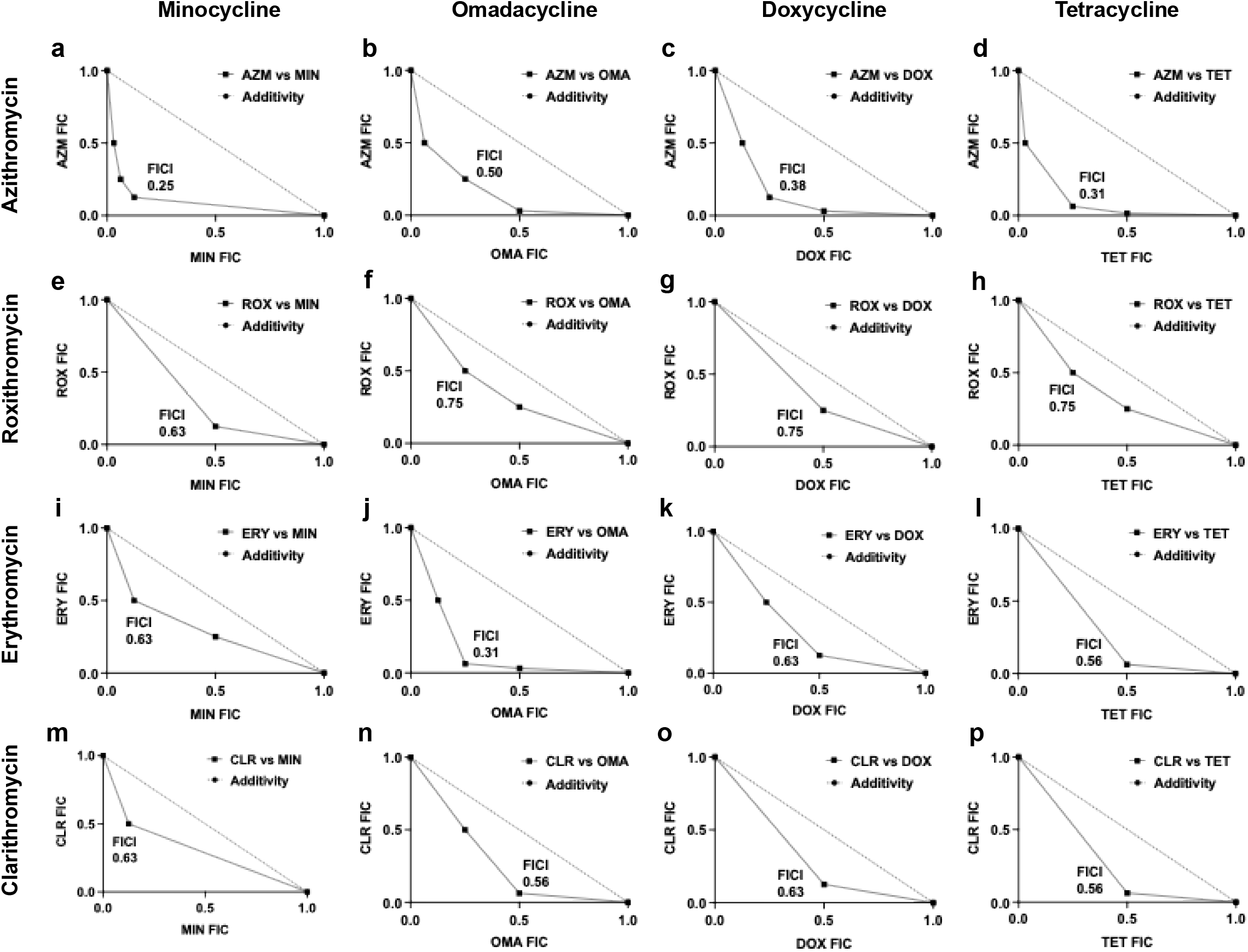
AZM combined with any tetracycline-class antibiotic enables potent dual-translation inhibition against AB5075 in CA-MHB. FIC assays of different macrolide and tetracycline combinations were performed against AB5075 in CA-MHB. In addition to AZM *(a - d)*, macrolide antibiotics included ROX (*e - h)*, ERY (*i - l)*, and CLR *(m - p)*. Each macrolide was assayed with a given tetracycline antibiotic, which consisted of MIN, OMA, DOX, and TET. All experiments were completed in triplicates, with a representative plot shown. FIC values are relative to 1.0, which is set as the MIC for each antibiotic in monotherapy. A synergistic relationship between the antibiotics is indicated when the FICI ≤ 0.5.

### AZM Exhibits Greater Kinetic-Based Translational Inhibition than Other Macrolides

The unique ability of AZM to drive synergy regardless of the tetracycline led us to hypothesize that differences in macrolide translation inhibitory potential dictate the strength of the synergistic interaction. We examined the inhibitory potential of each antibiotic using our established *A. baumannii* translation activity assays Translational activity is measured in these assays by utilizing a strain of MDR *A. baumannii* that constitutively expresses both the luciferase enzyme and luciferin substrate from the *luxCDBAE* luciferase cassette (AB5075-*luxCDBAE*). In this strain, inhibition of translation is associated with diminished luminescence. The inhibitory potential of each antibiotic was assessed through treating AB5075-*luxCDBAE* cultures with fractions of their individual MICs. A ratio of luminescence signal to optical density (OD_600_) was calculated to account for differences in signals due to bacterial concentrations. In monotherapy, macrolides were slow to inhibit translation but had sustained activity over time. AZM was the most potent antibiotic among the macrolides, significantly inhibiting translation at every concentration examined (1/4 MIC to 1/32 MIC) (Fig. 3 a-d). In contrast, ROX, ERY, and CLR only showed significant translation inhibition at 1/8 MIC fractions and above (Fig. 3 a-b). Unlike the macrolides’ delayed inhibitory response, the tetracycline drugs quickly inhibited translation at most sub-inhibitory concentrations. However, this inhibition was short-lived, as translational activity resumed after ∼7 hours. Except for the 1/4 MIC fractions for MIN and TET, no clear difference could be established from each tetracycline MIC fraction with respect to the control, supporting our hypothesis that tetracycline selection plays a minor role in influencing synergy (Fig. 3 e-h). These findings demonstrate that AZM is the most potent macrolide inhibitor of translation, while the tetracyclines inhibit translation similarly.

**Figure 3.**
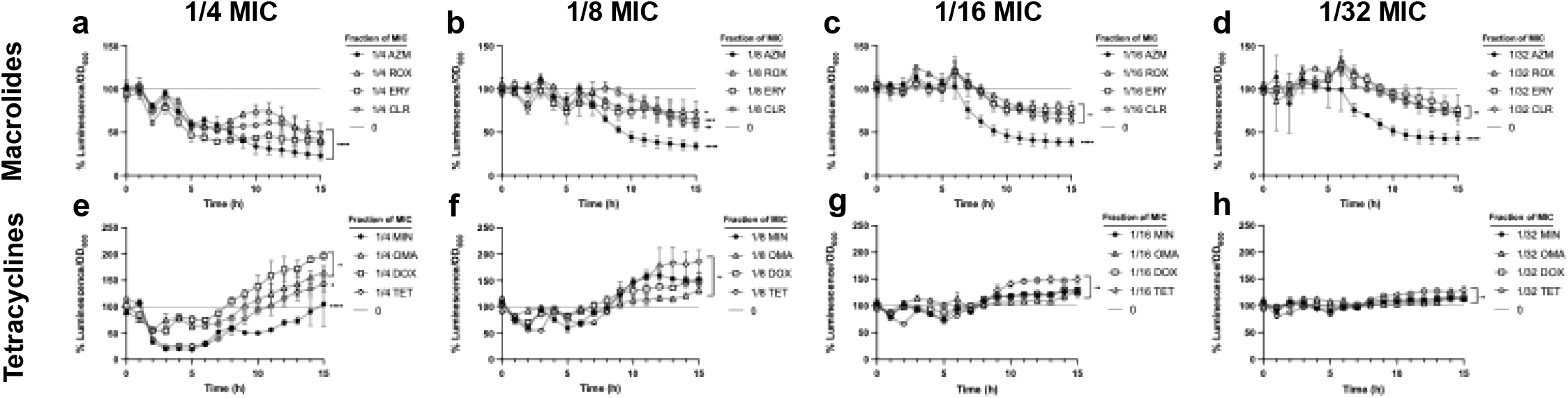
Sub-inhibitory AZM, but not MIN, has enhanced translation inhibition compared to other members of its antibiotic class. Monotherapy translation inhibition assays were performed for each macrolide (*a - d*) and tetracycline (*e – h*) antibiotic at fractions of the MIC and against the AB5075-*luxCDABE* strain in CA-MHB. Translational activity was assessed on a cellular level independent of the bacterial concentration by calculating the ratio between the luminescence and cellular density, as measured by OD_600_. Assays were run for 15 hours with luminescence and OD_600_ values recorded at 30-minute intervals. All luminescence/OD_600_ ratios were graphed as a percentage of untreated AB5075-*luxCDABE* growth. All experiments were conducted in triplicates. Error bars were calculated by using the standard deviation for each point. Statistical significance was calculated using a one-way ANOVA with * ≤0.0500–0.0100, ** ≤0.0100– 0.0010, *** ≤0.0010–0.0001, and **** ≤0.0001.

### Dual-Therapy Translation Inhibition Assays Underlie AZM Efficacy in Kinetic-Based Synergism

After identifying AZM as the most potent macrolide inhibitor of translation, we next examined the combined effect of macrolide-tetracycline pairings. We conducted dual-therapeutic translation inhibition assays for each combination at 1/4 MIC fractions against AB5075-*luxCDBAE*. In agreement with our FIC data (Fig. 2), all AZM-tetracycline combinations were found to significantly inhibit translation (Fig. 4-5). In addition to AZM, ERY + OMA, also significantly inhibited translation, through to a lesser extent than any of the AZM combinations. Contrary to the synergistic combinations, most non-synergistic pairings did not display statistically significant inhibition compared to the untreated control, except for the ROX + TET, ERY + MIN, and ERY + TET pairings (Fig. 2.6-2.7). The similarity in results between dual-therapeutic assays where the macrolide is the same, but the tetracycline differs, supports the idea that the type of tetracycline is less influential in determining synergy than the type of macrolide. Furthermore, the strong translation inhibition in all AZM combinations, but not for any other macrolide, highlight AZM as the most effective macrolide in facilitating synergy due to its robust, long-lasting inhibition complementing the potent but short-lived action of the paired tetracycline.

**Figure 4.**
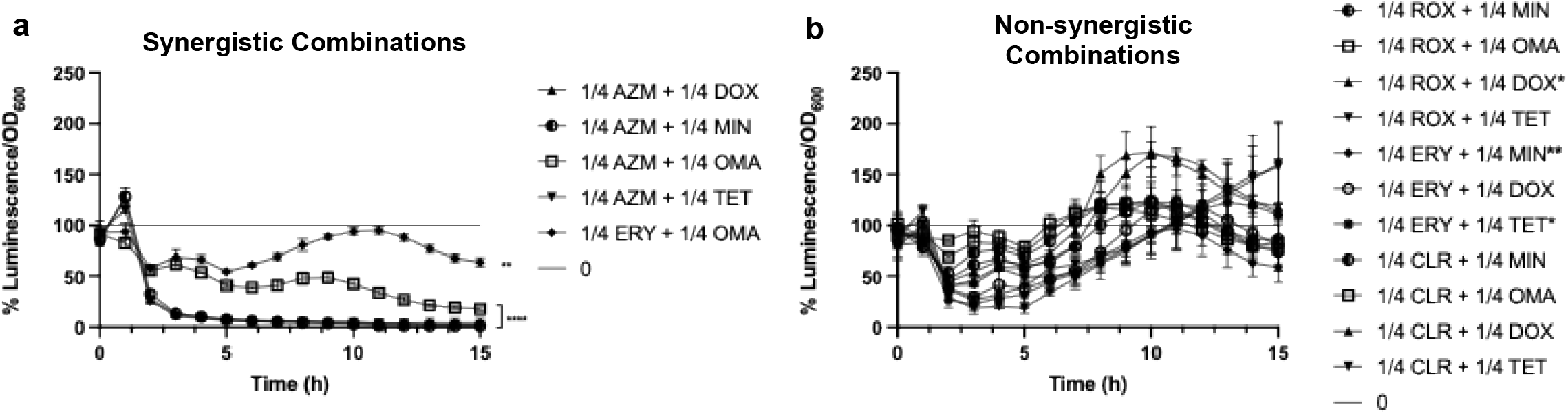
Synergistic macrolide-tetracycline combination therapies displayed robust translation inhibition unlike non-synergistic combinations. Dual antibiotic translation inhibition assays were conducted for each macrolide-tetracycline pair at 1/4 MIC fractions against the AB5075-*luxCDABE* strain in CA-MHB. Translational activity curves of synergistic dual-therapies (a) and non-synergistic dual-therapies (b) were plotted on separate graphs. Translational activity was assessed on a cellular level independent of the bacterial concentration by calculating the ratio between the luminescence and cellular density, as measured by OD_600_. Assays were run for 15 hours with luminescence and OD_600_ values recorded at 30-minute intervals. All luminescence/OD_600_ ratios were graphed as a percentage of untreated AB5075-*luxCDABE* growth. All experiments were conducted in triplicates. Error bars were calculated by using the standard deviation for each point. Statistical significance was calculated using a one-way ANOVA with * ≤0.0500–0.0100, ** ≤0.0100–0.0010, *** ≤0.0010–0.0001, and **** ≤0.0001.

**Figure 5.**
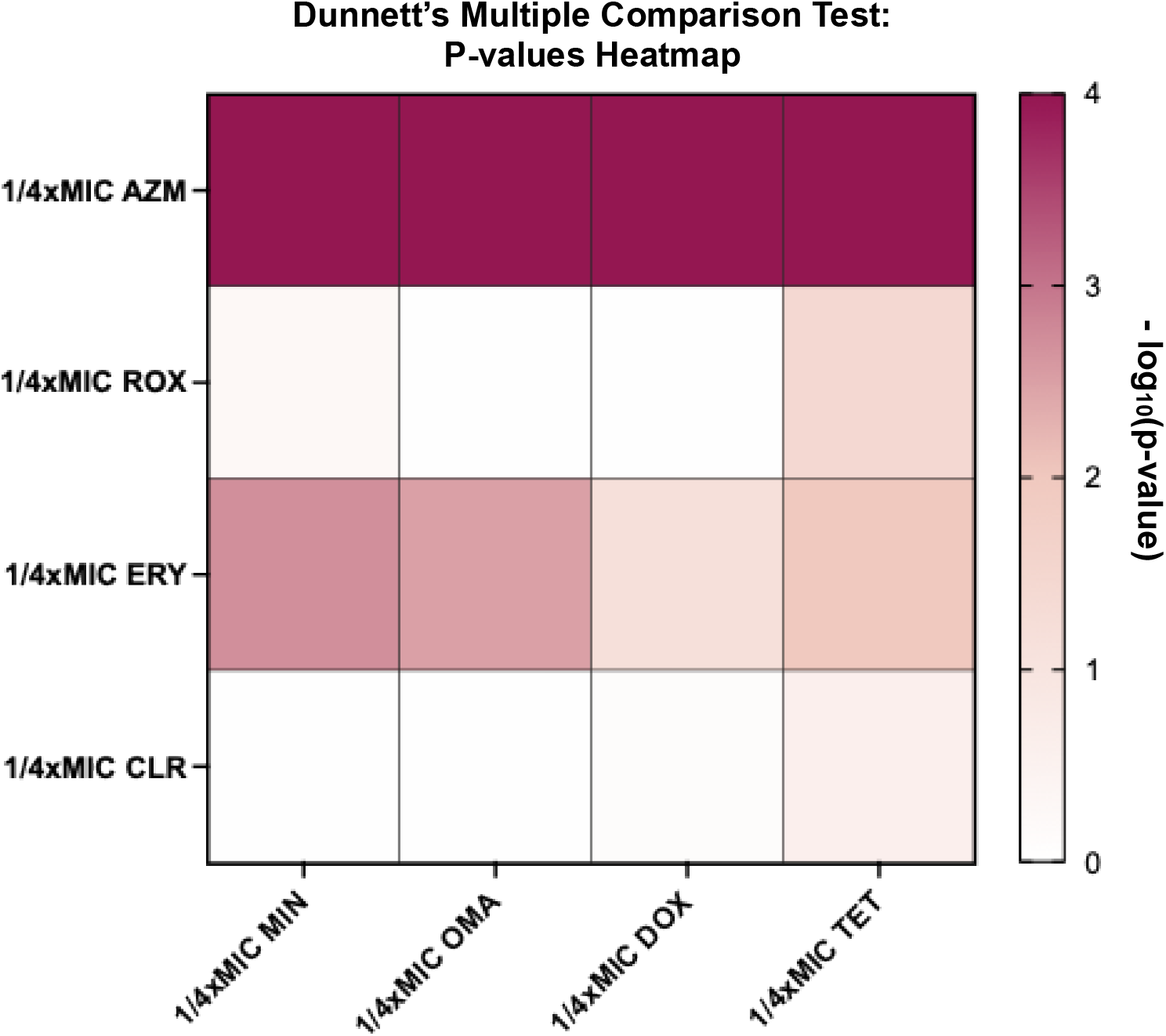
AZM combination therapies display significantly greater inhibition than any other combination therapy. A Dunnett’s comparison test was performed to evaluate the significant difference in translational inhibitory capacity between each 1/4 MIC combination therapy and an untreated control. P-values were logarithmically transformed and scaled from 0 to 4 on a 4×4 heat map, ranging respectively from no significance to high significance. 1/4 Macrolides are organized by rows, and 1/4 tetracyclines are organized by columns. Statistical significance was calculated using a one-way ANOVA with * ≤0.0500–0.0100, ** ≤0.0100–0.0010, *** ≤0.0010–0.0001, and **** ≤0.0001.

### AZM-MIN Synergy can overcome resistance to MIN monotherapy

Having established that AZM drives synergistic macrolide-tetracycline combinations, we concluded our investigation by assessing the efficacy of AZM-MIN synergy against 41 MDR *A. baumannii* strains from the CDC-FDA AR isolate bank (17). The MIC_90_ of MIN and AZM was first determined for each strain in both CA-MHB and RPMI+ (Table S1). Unlike with earlier results in Table 1, most of the AR strains were resistant to MIN (36 strains), as defined by the clinical resistance breakpoints (MIC_90_ of MIN ≥ 16 µg/ml) set by the CLSI guidelines (21). MIC_90_ values for MIN ranged from 8 - 256 µg/ml in CA-MHB and >256 µg/ml in RPMI+ (Table S1). The MIC_90_ values for AZM in CA-MHB were high in 38/41 strains (≥512 µg/ml), with only three of the 41 having an MIC90 <512 µg/ml. In RPMI+, however, AZM activity noticeably improved against some of the isolates, with nine of the 41 strains having a MIC_90_ < 8 µg/ml (Table S1).

We next conducted a series of FIC assays to assess if AZM and MIN synergizes against any of the 41 strains in CA-MHB. FICI values for AZM-MIN combination therapies against all 41 strains are listed in Table S1. Eight of the 41 strains were revealed to be susceptible to the combination therapy (Table 3). We then performed another series of FIC assays against only the susceptible strains in RPMI+, assessing for MIN-AZM synergism. AZM-MIN was enhanced in RPMI+ as evidenced by the reduced FICIs in seven out of the eight susceptible strains (Table 3). Furthermore, of the eight susceptible strains, AZM-MIN combination therapy lowered the MIN concentration to that of below the CLSI clinical breakpoint for four strains in CA-MHB. (Table 4)

**Table 3.** MIC_90_ (µg/ml) & FICI of select *A. baumannii* strains from the CDC *A. baumannii* AR bank containing MDR clinical isolates.

|  | MIC<sub>90</sub> in CA-MHB |  | MIC<sub>90</sub> in RPMI+ |  | FICI against AZM-MIN Combination |  |
| --- | --- | --- | --- | --- | --- | --- |
|  | MIN | AZM | MIN | AZM | CA-MHB | RPMI+ |
| *Acinetobacter baumannii* |  |  |  |  |  |  |
| -AR0273 | 64 | 128 | >256 | 1 | 0.375 | 0.15625 |
| -AR0289 | 32 | >512 | >256 | 0.5 | 0.3125 | 0.15625 |
| -AR0292 | 256 | >512 | >256 | 128 | 0.3125 | 0.125 |
| -AR0293 | 16 | 128 | >256 | 1 | 0.28125 | 0.3125 |
| -AR0294 | 16 | >512 | >256 | 0.5 | 0.5 | 0.140625 |
| -AR0304 | 32 | >512 | >256 | 0.5 | 0.375 | 0.1875 |
| -AR0305 | 2 | >512 | >256 | 1 | 0.5 | 0.125 |
| -AR0313 | 64 | >512 | >256 | 4 | 0.5 | 0.09375 |

**Table 4.** FICs and inhibitory MIN & AZM concentrations (µg/ml) of combination therapies against select AR isolates in CA-MHB.

|  | FICs in CA-MHB |  | Inhibitory Concentrations in CA-MHB MIN-AZM Synergistic Therapy |  | Meets MIN CLSI Susceptibility Breakpoint (≤ 4 µg/ml) |
| --- | --- | --- | --- | --- | --- |
|  | MIN | AZM | MIN | AZM |  |
| *Acinetobacter baumannii* |  |  |  |  |  |
| - AR0273 | 0.25 | 0.125 | 16 | 16 | N |
| - AR0289 | 0.0625 | 0.25 | 2 | 256 | Y |
| - AR0292 | 0.25 | 0.0625 | 64 | 64 | N |
| - AR0293 | 0.03125 | 0.25 | 0.5 | 32 | Y |
| - AR0294 | 0.25 | 0.25 | 4 | 256 | Y |
| - AR0304 | 0.125 | 0.25 | 4 | 256 | Y |
| - AR0305 | 0.25 | 0.25 | 0.5 | 32 | N/A |
| - AR0313 | 0.25 | 0.25 | 16 | 256 | N |

## Discussion

The rise of MDR *A. baumannii* continues to challenge clinical treatment paradigms. In prior studies, AZM-MIN synergism was established as an effective treatment against MDR *A. baumannii* in vitro but no key drivers of synergism were identified. Furthermore, whether AZM-MIN synergism was strain-or if its potency correlated with enhanced translation inhibition were also unclear (10). In this study, we identify AZM, a macrolide not conventionally used for Gram-negative infections, as a critical synergistic partner when combined with tetracycline class antibiotics, especially MIN (Fig. 2b, FICI = 0.25). Furthermore, our established translation inhibition assays characterized AZM as a potent inhibitor with greater longitudinal activity in sub-inhibitory concentrations than the other selected macrolides. These findings were supplemented by our dual-antibiotic inhibition assays, where AZM paired combination therapies demonstrated significantly enhanced inhibition, unlike most other macrolide-tetracycline combinations (Fig. 4). Application of AZM-MIN combination therapy against each of the 41 MDR *A. baumannii* strains from the AR isolate bank revealed AZM-MIN synergism can occur against 8 of those strains in either CA-MHB or RPMI+ media. In 4 of the selected 8 strains, AZM-MIN synergism reduced the therapeutic MIN concentration in CA-MHB from at or above the CLSI resistance clinical breakpoint (16 µg/ml) to at or below the CLSI susceptibility clinical break point of 4 µg/ml (Table 4) (21). In turn, our studies show that AZM-MIN synergistic therapy may potentially resensitize resistant *A. baumannii* strains to MIN (22).

Our findings offer several practical advantages. First, AZM-MIN synergy reduced the effective concentration of MIN below the CLSI resistance breakpoint in 4 out of 8 MDR isolates that responded to the combination (21). This suggests that even strains resistant to MIN monotherapy could become treatable through combination therapy, expanding the clinical utility of existing drugs. Second, as synergy was observed with all tetracyclines tested (though most robustly with MIN), AZM can be paired with alternative tetracyclines like OMA in cases where resistance mechanisms (e.g., *tetB*) limit MIN efficacy (Fig. 2b, FICI = 0.5) (23, 24). This flexibility allows treatment strategies to be tailored to specific resistance profiles, while minimizing selection pressure for MIN resistance which is a critical benefit in preserving the longevity of antibiotics. Beyond *A. baumannii*, the implications of our findings extend to other Gram-negative pathogens. Given the growing interest in dual-translation inhibitors and non-conventional combination therapies, AZM-based therapies could be repurposed to improve outcomes in infections caused by *Pseudomonas aeruginosa* or *Klebsiella pneumoniae*.

Our findings also suggest that bicarbonate enhances the efficacy of AZM and its synergy with tetracyclines. AZM MIC_90_ values were consistently lower in RPMI+ (which contains bicarbonate) compared to CA-MHB (Tables 1 and S1), and AZM-MIN synergy was notably stronger in RPMI+ as indicated by reduced FICIs (Figure 1, Table 3). Of the 8 AZM-MIN-susceptible MDR strains, 7 displayed low AZM MIC_90_ (≤ 4 µg/mL) in RPMI+, supporting a strong link between bicarbonate-enhanced AZM activity and synergy. Our results aligned with prior reports that bicarbonate can inhibit bacterial growth of cystic fibrosis pathogens and potentiate macrolide activity via disruption of membrane potential (25, 26). These findings underscore the potential of leveraging host-like bicarbonate conditions to improve AZM-based therapies, especially in infections like cystic fibrosis or pneumonia where bicarbonate is physiologically relevant. Understanding how bicarbonate modulates antibiotic activity could lead to optimized treatment protocols or even adjunctive therapies that manipulate host environments to boost drug efficacy. Future studies should directly test AZM-MIN combinations in bicarbonate rich and deficient conditions and dissect the molecular mechanisms underlying bicarbonate-mediated potentiation in *A. baumannii* and other Gram-negative pathogens.

Lastly, the kinetic model of sequential translation inhibition presented here offers a new approach for antimicrobial synergy. While most combination therapies rely on distinct mechanisms of action, our approach uses two translation inhibitors whose complementary kinetics expand the window of protein synthesis disruption. This synergy could represent a broader strategy for designing rational antibiotic combinations that exploit drug pharmacodynamics rather than simply combining distinct molecular targets.

In summary, our study provides compelling mechanistic and translational evidence supporting the use of AZM-tetracycline combinations against MDR *A. baumannii*. AZM is revealed as a potent enabler of synergy, capable of restoring MIN susceptibility in resistant isolates. These findings open new avenues for reviving the clinical utility of existing antibiotics and developing kinetic-based combination therapies tailored to MDR infections. Future in vivo and clinical investigations will be crucial to fully realize the potential of AZM-based therapies in Gram-negative bacterial infections.

## Contributors

Syed Raza, Suman Tiwari, and Dr. Dillon had full access to all data in the study and take responsibility for the integrity of the data and the accuracy of data analysis.

Study concept and design: Raza, Tiwari, Dillon.

Acquisition, analysis or interpretation of data: Raza, Tiwari, Dillon.

Laboratory experiments: Raza.

Drafting of manuscript: Raza, Tiwari, Dillon.

Administrative, technical or material support: Tiwari, Dillon.

Statistical analysis: Raza.

Critical revision of the manuscript for important intellectual content: Raza, Tiwari, Dillon

## Data Sharing Statement

## Supplementary Data

**Figure S1.**
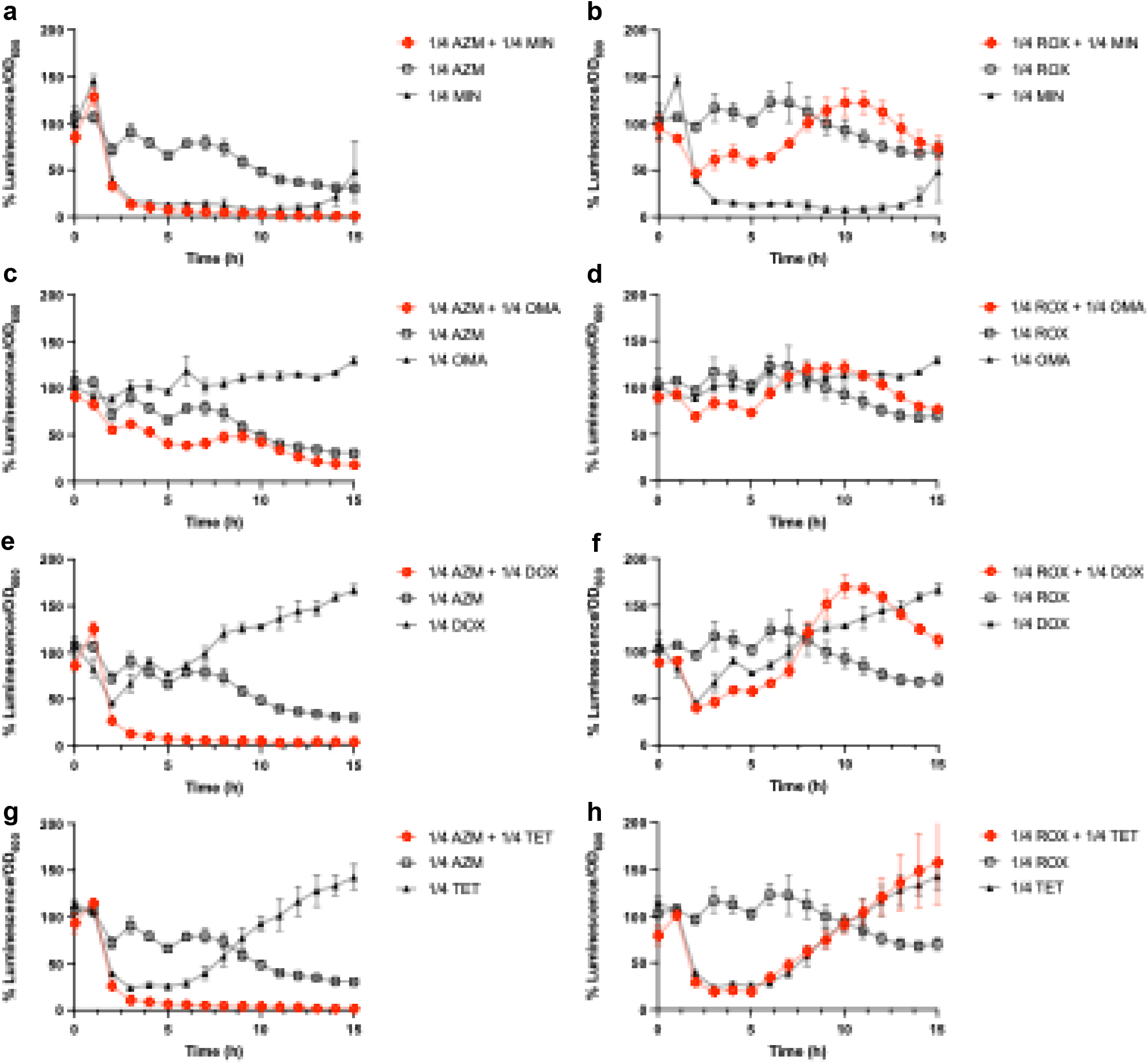
AZM-paired combination therapies show significantly greater inhibitory activity than either macrolide or tetracycline constituent in monotherapy. 1/4 MIC monotherapy translation inhibition assays were plotted on the same graph with the corresponding 1/4 dual antibiotic translation inhibition assays. AZM combination therapies (*a – d)* demonstrated significantly greater translation inhibition relative to their constituent 1/4 MIC monotherapy assays than the ROX (*e – h)* combination therapy. Translational activity was assessed on a cellular level independent of the bacterial concentration by calculating the ratio between the luminescence and cellular density, as measured by OD_600_. Assays were run for 15 hours with luminescence and OD_600_ values recorded at 30-minute intervals. All luminescence/OD_600_ ratios were graphed as a percentage of untreated AB5075-*luxCDABE* growth. All experiments were conducted in triplicates. Error bars were calculated by using the standard deviation for each point. Statistical significance was calculated using a one-way ANOVA with * ≤0.0500–0.0100, ** ≤0.0100–0.0010, *** ≤0.0010–0.0001, and **** ≤0.0001.

**Figure S2.**
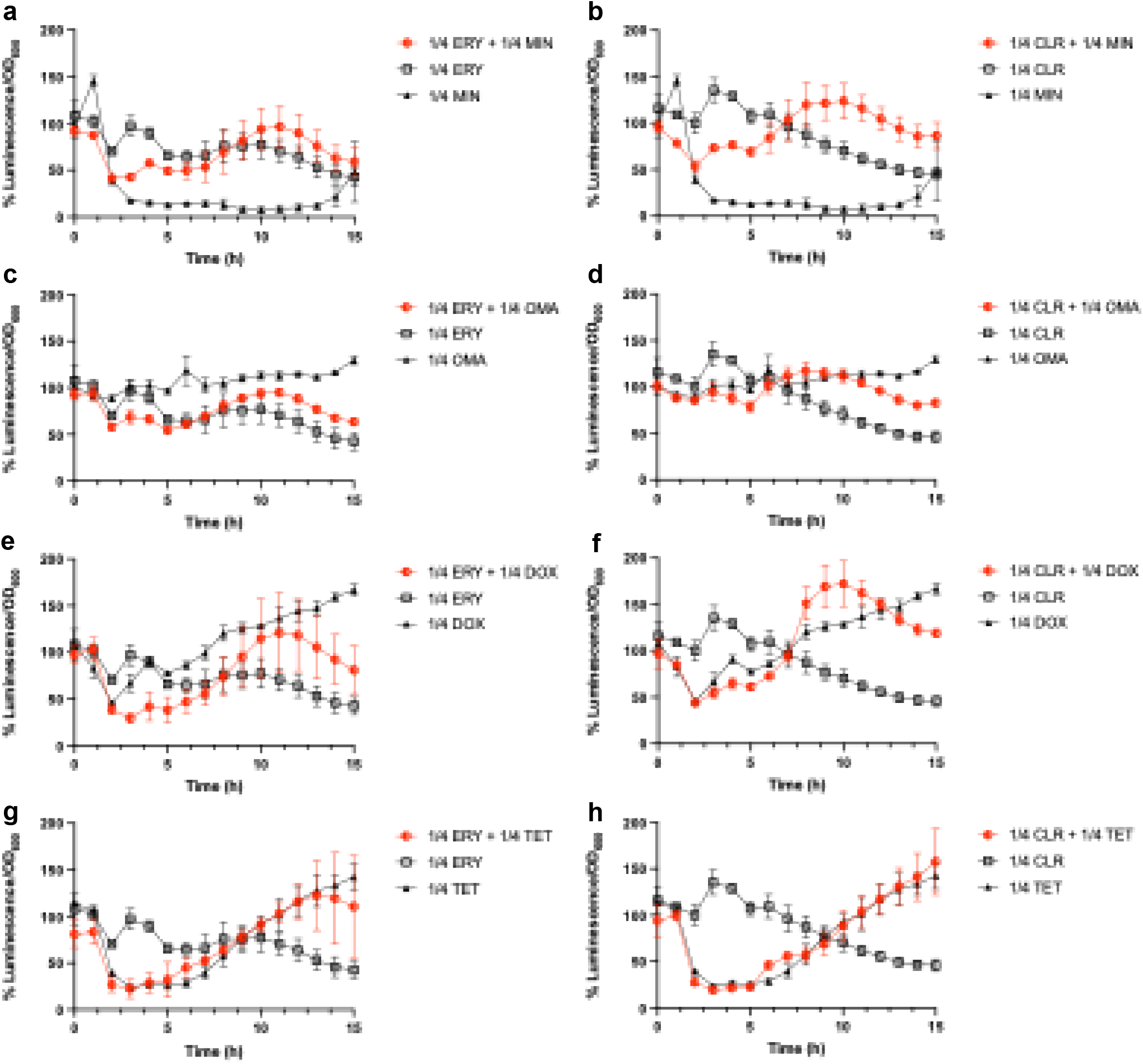
Similar to ROX-paired combinations and unlike AZM-paired combinations, ERY and CLR-paired combinations to do show significantly greater inhibitory activity than their constituent monotherapies. 1/4 MIC monotherapy translation inhibition assays were plotted on the same graph with the corresponding 1/4 dual antibiotic translation inhibition assays. ERY (*a – d*) and CLR (*e – h*) combination therapies do not display significantly greater translation inhibition relative to their constituent 1/4 MIC monotherapy assays, unlike AZM combination therapies. Translational activity was assessed on a cellular level independent of the bacterial concentration by calculating the ratio between the luminescence and cellular density, as measured by OD_600_. Assays were run for 15 hours with luminescence and OD_600_ values recorded at 30-minute intervals. All luminescence/OD_600_ ratios were graphed as a percentage of untreated AB5075-*luxCDABE* growth. All experiments were conducted in triplicates. Error bars were calculated by using the standard deviation for each point. Statistical significance was calculated using a one-way ANOVA with * ≤0.0500–0.0100, ** ≤0.0100–0.0010, *** ≤0.0010–0.0001, and **** ≤0.0001.

**Figure S3.**
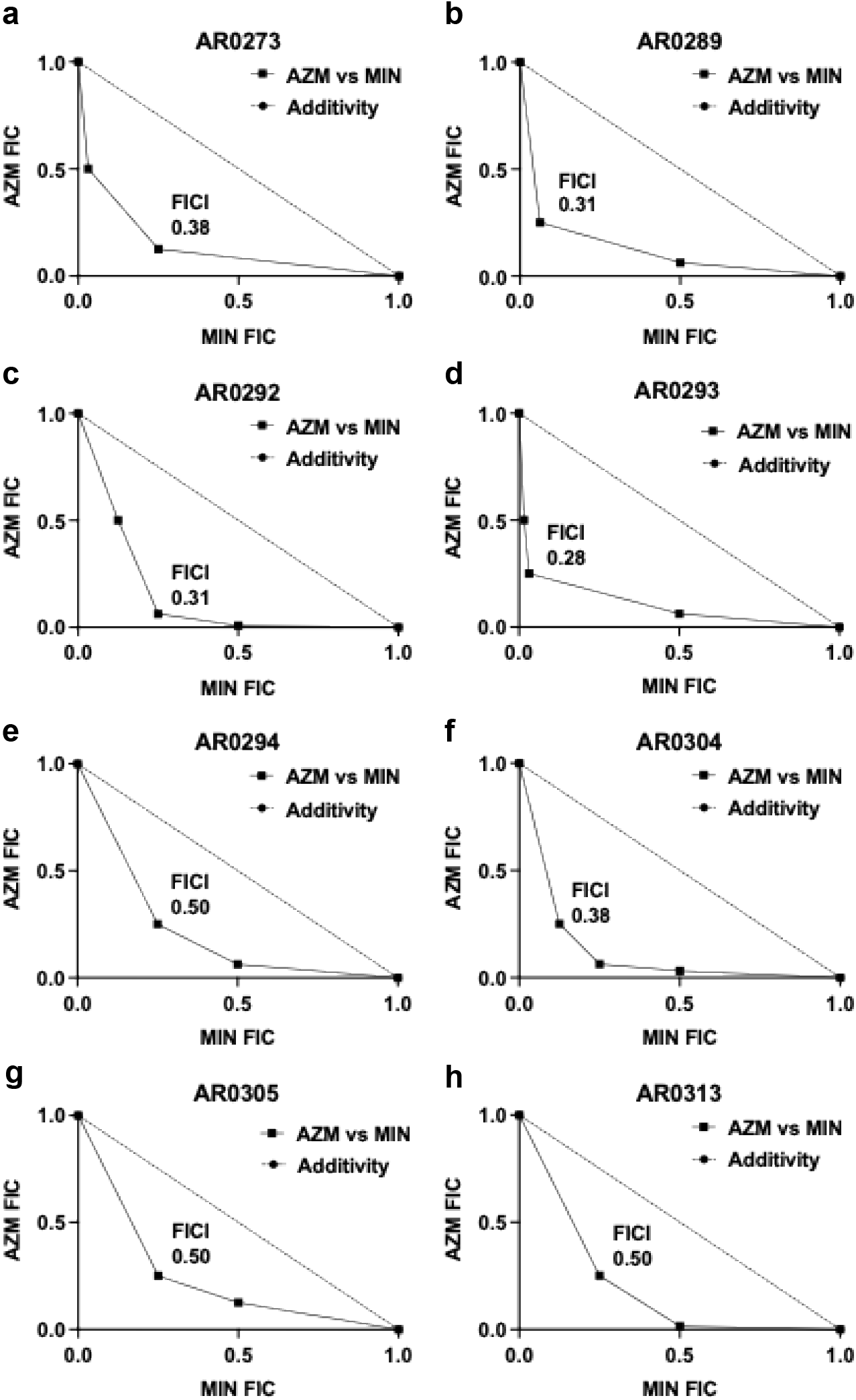
8 AR *A. baumannii* isolates are susceptible to AZM-MIN synergy in CA-MHB. 41 MDR *A. baumannii* strains that are part of the CDC-FDR AR isolate bank were tested for AZM-MIN synergy through our FIC assays. Of the 41 strains, AZM-MIN synergized against 8 of those strains in CA-MHB (*a –h)*. All experiments were completed in triplicates, with a representative plot shown. FIC values are relative to 1.0, which is set as the MIC for each antibiotic in monotherapy. A synergistic relationship between the antibiotics is indicated when FICI ≤ 0.5.

**Figure S4.**
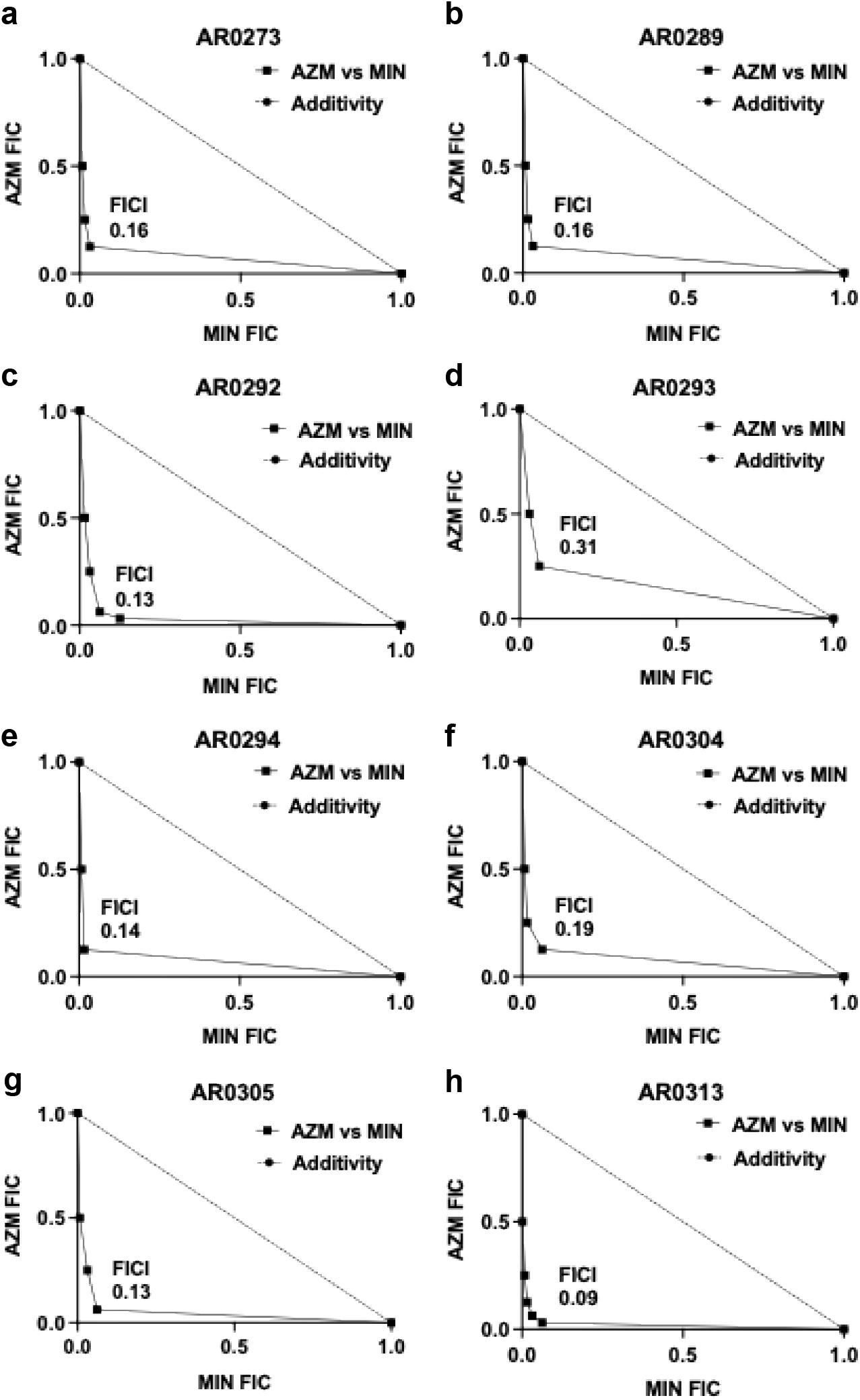
8 AR *A. baumannii* isolates are susceptible to AZM-MIN synergy in RPMI+. FIC testing was repeated in RPMI+ but only for the 8 strains that showed synergism in CA-MHB. AZM-MIN synergized against the 8/8 susceptible strains in RPMI+ (*a – h)*, yielding potentiated synergism than in CA-MHB. All experiments were completed in triplicates, with a representative plot shown. FIC values are relative to 1.0, which is set as the MIC for each antibiotic in monotherapy. A synergistic relationship between the antibiotics is indicated when FICI ≤ 0.5.

**Table S1.**
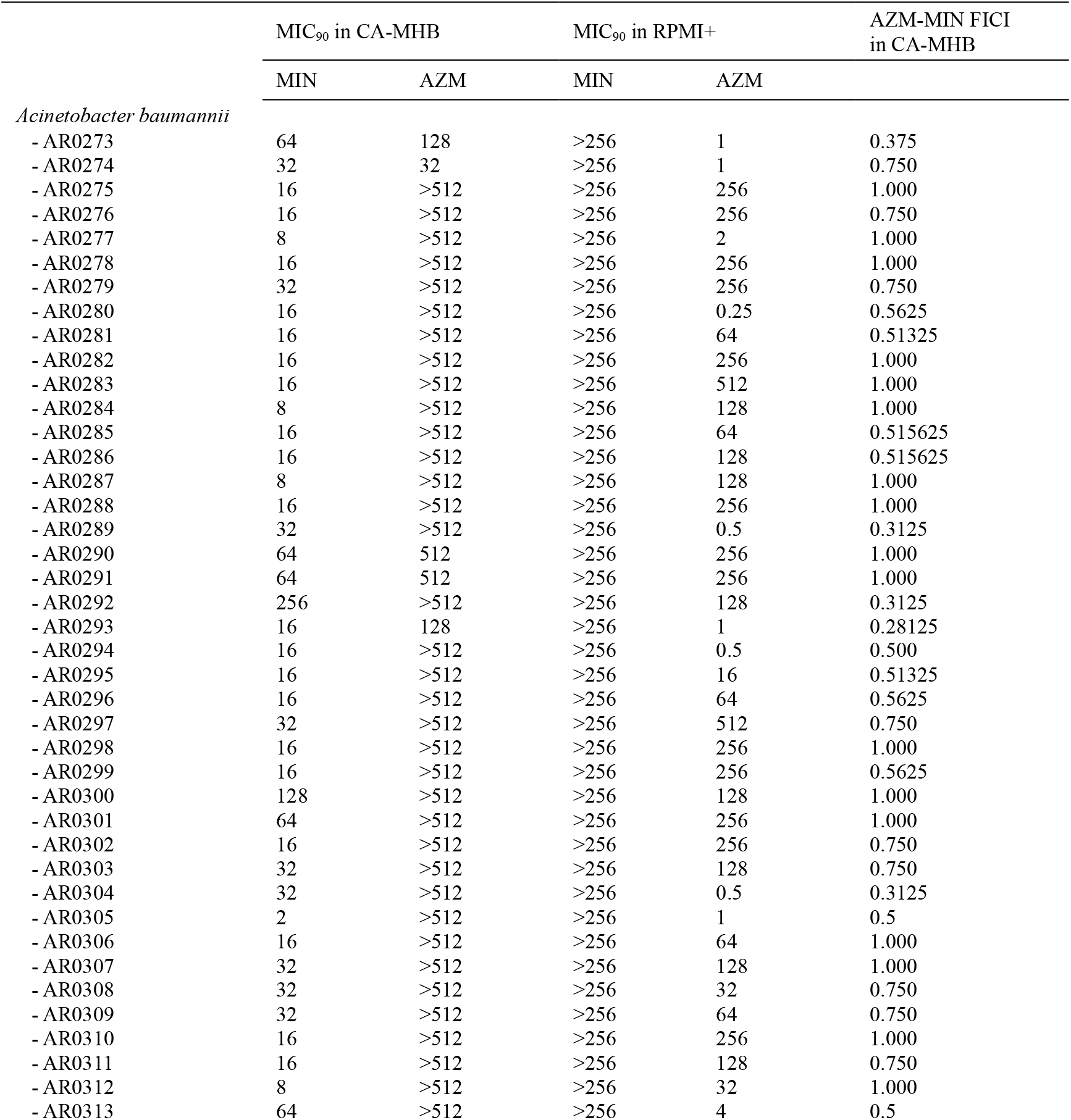
MIC_90_ (µg/mL) & FICI of *A. baumannii* strains from the CDC *A. baumannii* AR bank containing MDR clinical isolates.

|  | MIC<sub>90</sub> in CA-MHB |  | MIC<sub>90</sub> in RPMI+ |  | AZM-MIN FICI<br>in CA-MHB |
| --- | --- | --- | --- | --- | --- |
|  | MIN | AZM | MIN | AZM |  |
| *Acinetobacter baumannii* |  |  |  |  |  |
| - AR0273 | 64 | 128 | >256 | 1 | 0.375 |
| - AR0274 | 32 | 32 | >256 | 1 | 0.750 |
| - AR0275 | 16 | >512 | >256 | 256 | 1.000 |
| - AR0276 | 16 | >512 | >256 | 256 | 0.750 |
| - AR0277 | 8 | >512 | >256 | 2 | 1.000 |
| - AR0278 | 16 | >512 | >256 | 256 | 1.000 |
| - AR0279 | 32 | >512 | >256 | 256 | 0.750 |
| - AR0280 | 16 | >512 | >256 | 0.25 | 0.5625 |
| - AR0281 | 16 | >512 | >256 | 64 | 0.51325 |
| - AR0282 | 16 | >512 | >256 | 256 | 1.000 |
| - AR0283 | 16 | >512 | >256 | 512 | 1.000 |
| - AR0284 | 8 | >512 | >256 | 128 | 1.000 |
| - AR0285 | 16 | >512 | >256 | 64 | 0.515625 |
| - AR0286 | 16 | >512 | >256 | 128 | 0.515625 |
| - AR0287 | 8 | >512 | >256 | 128 | 1.000 |
| - AR0288 | 16 | >512 | >256 | 256 | 1.000 |
| - AR0289 | 32 | >512 | >256 | 0.5 | 0.3125 |
| - AR0290 | 64 | 512 | >256 | 256 | 1.000 |
| - AR0291 | 64 | 512 | >256 | 256 | 1.000 |
| - AR0292 | 256 | >512 | >256 | 128 | 0.3125 |
| - AR0293 | 16 | 128 | >256 | 1 | 0.28125 |
| - AR0294 | 16 | >512 | >256 | 0.5 | 0.500 |
| - AR0295 | 16 | >512 | >256 | 16 | 0.51325 |
| - AR0296 | 16 | >512 | >256 | 64 | 0.5625 |
| - AR0297 | 32 | >512 | >256 | 512 | 0.750 |
| - AR0298 | 16 | >512 | >256 | 256 | 1.000 |
| - AR0299 | 16 | >512 | >256 | 256 | 0.5625 |
| - AR0300 | 128 | >512 | >256 | 128 | 1.000 |
| - AR0301 | 64 | >512 | >256 | 256 | 1.000 |
| - AR0302 | 16 | >512 | >256 | 256 | 0.750 |
| - AR0303 | 32 | >512 | >256 | 128 | 0.750 |
| - AR0304 | 32 | >512 | >256 | 0.5 | 0.3125 |
| - AR0305 | 2 | >512 | >256 | 1 | 0.5 |
| - AR0306 | 16 | >512 | >256 | 64 | 1.000 |
| - AR0307 | 32 | >512 | >256 | 128 | 1.000 |
| - AR0308 | 32 | >512 | >256 | 32 | 0.750 |
| - AR0309 | 32 | >512 | >256 | 64 | 0.750 |
| - AR0310 | 16 | >512 | >256 | 256 | 1.000 |
| - AR0311 | 16 | >512 | >256 | 128 | 0.750 |
| - AR0312 | 8 | >512 | >256 | 32 | 1.000 |
| - AR0313 | 64 | >512 | >256 | 4 | 0.5 |

